# Simplified, Shear Induced Generation of Double Emulsions for Robust Compartmentalization during Single Genome Analysis

**DOI:** 10.1101/2021.11.22.469484

**Authors:** Thomas W. Cowell, Andrew Dobria, Hee-Sun Han

## Abstract

Drop microfluidics has driven innovations for high throughput, low input analysis techniques such as single-cell RNA-seq. However, the instability of single emulsion (SE) drops occasionally causes significant merging during drop processing, limiting most applications to single-step reactions in drops. Here, we show that double emulsion (DE) drops address this critical limitation and completely prevent content mixing, which is essential for single entity analysis. DEs show excellent stability during thermal cycling. More importantly, DEs undergo rupture into the continuous phase instead of merging, preventing content mixing and eliminating unstable drops from the downstream analysis. Due to the lack of drop merging, the monodispersity of drops is maintained throughout a workflow, enabling the deterministic manipulation of drops downstream. We also developed a simple, one-layer fabrication method for DE drop makers. This design is powerful as it allows robust production of single-core DEs at a wide range of flow rates and better control over the shell thickness, both of which have been significant limitations of conventional two-layer devices. This approach makes the fabrication of DE devices much more accessible, facilitating its broader adoption. Finally, we show that DE droplets effectively maintain the compartmentalization of single virus genomes during PCR-based amplification and barcoding, while SEs mixed contents due to merging. With their resistance to content mixing, DE drops have key advantages for multistep reactions in drops, which is limited in SEs due to merging and content mixing.

## 1. Introduction

Droplet microfluidics has established itself for studying biological and biochemical systems with high throughput and low cost.^1–4^ By forming millions of near-identical water in oil droplets, samples are partitioned enabling in-drop directed evolution, high throughput screening, and single-molecule or single-cell analysis.^5–7^ These foundational techniques have relied upon the efficient compartmentalization enabled by monodisperse SE droplets. However, stabilizing SEs remains a persistent challenge.^7^ Despite the significant effort that has been devoted to developing surfactants that stabilize SE droplets during incubation, thermal cycling, reinjection, and other drop manipulations, drop merging cannot be fully avoided.^8–10^ SE systems are highly sensitive, with drop coalescence varying with drop size^11,12^ aqueous phase composition,^13–15^ surfactant composition,^8,9^ thermal gradients,^16,17^ and shear stress^18^. The extent of drop merging can also vary significantly between experimental replicates as shown in **Figure S1**. Maintaining a monodisperse SE throughout a multistep droplet workflow is especially difficult as disturbances to the drop size distribution compound with each operation.^7,19^ Drop merging can result in a loss of single entity resolution as contents mix, or uneven sequencing reads as larger drops amplify with higher yield. Moreover, drop-by-drop manipulations require monodisperse inputs to stablize flow. Consequently, the commercialized microfluidic techniques that use SE droplets are limited to at most one drop generation and one incubation or thermal cycling step before analysis or pooling in bulk.^20,21^ This stability-induced limitation dramatically narrows the scope of possible applications for SE microfluidics.

We believe that DE droplets can provide a good solution for this long-standing problem. Since DEs are composed of aqueous compartments partitioned from the continuous aqueous phase by an immiscible oil shell, we hypothesized that their coalescence behavior would be distinct from SEs. Instead of drop merging, which causes SE compartments to combine and mix their contents, DE drops undergo drop rupture, which releases the aqueous contents into the continuous phase effectively removing the coalesced material from the workflow and preserving the monodispersity of the emulsion despite coalescence. This unique coalescence behavior is well suited for lengthy or multistep protocols, as ruptured material is removed from downstream steps leaving behind only the intact emulsion for subsequent drop manipulations. Moreover, recent demonstrations have shown DE compatibility with sorting by FACS instruments.^22–24^ Sorting by commercial flow cytometers eliminates the need for complex optical and electronic equipment and the associated expertise which is required to sort SEs.^4,7,25^ This combination of factors, positions DEs as a promising modality in the proliferation of droplet-based high throughput assays and screening. In this paper, we test our hypothesis that DE drops effectively eliminate drop merging and content mixing and highlight the strength of DE-based approaches for single-cell analysis over traditional SE methods. We also significantly lower the barrier to access the DE technologies by developing a simple and robust drop maker.

The primary unresolved impediments to the wider use of DE techniques for high-throughput screening and single-cell profiling is the complexity of fabrication and operation of DE droplet generators. Traditional DE drop making devices use either a 3D nested channel geometry and/or surface wetting control to achieve water-in-oil-in-water emulsification.^26–29^ 3-D nested channel geometries position an incoming stream of water-in-oil droplets within a surrounding aqueous flow ensuring that the formation of the DE drop is correct, independent of the surface wetting of the channels. This concentric channel architecture is most commonly achieved by aligning glass capillaries,^30–32^ 3D printing,^33–35^ or by multi-layer soft lithography.^36,37^ These fabrication methods, however, require complex equipment, expensive facilities, and/or technical expertise to perform.^38–40^ Further, glass capillary fabrication is limited to larger channel dimensions and faces greater device-to-device variation,^41^ which can make the production of the small DE drops that are suitable for flow cytometry sorting,^42^ more challenging. To avoid full 3-D fabrication, semi-planar DE drop makers have been designed to use surface wetting to control emulsification. These devices feature a planar lower channel surface but utilize a step increase in channel height at the second junction to assist spatial patterning of surface treatments and DE formation. For both device designs, drop formation at each junction must be synchronized to form quality DEs. Small changes in flow rates can result in improper loading of multiple cores into DE drops.^43^ As a result, the successful operation of these DE drop makers requires patient optimization.

When forming DEs in planar or semi-planar devices, the surface wetting of the channels must switch from hydrophobic to hydrophilic between the first and second junctions. Accordingly, spatially control of the surface wetting is required for DE formation using this approach.^44–46^ Previous efforts have demonstrated polymer deposition for the spatial patterning of the surface wetting of PDMS microchannels.^47,48^ Although effective, polymer deposition methods are laborious to perform as the in-flow of the polymer treatment solution must be carefully balanced by a second fluid that blocks the treatment solution from entering the untreated portions of the device. This flow balancing is typically aided by a semi-planar geometry where the device height is greater downstream of the second junction.^48,49^ The larger channel cross-section in the downstream portion of the device allows the hydrophilic polymer solution to flow in with less resistance than in the narrower portion of the device that is to remain hydrophobic. While effective, this semi-planar configuration still requires the alignment of photoresist or PDMS layers during fabrication, requiring technical expertise for fabrication.^50,51^ Plasma treatment was identified as a suitable alternative to apply temporary hydrophilic surface treatments to PDMS without flow balancing.^49,52,53^ Studies have shown spatial control of plasma-induced surface hydrophilization suitable for DE formation.^22-24,52^ Again, these devices utilize a semi-planar configuration to facilitate plasma diffusion into the larger channels of the hydrophilic portion of the device.

Here, we report the first fully planar DE drop maker to our knowledge, greatly enhancing the accessibility of DE systems, and we demonstrate how the unique coalescence behavior of DEs effectively maintains compartmentalization. This single-layer device reduces fabrication complexity to a minimum. Comparisons between 2-layer semi-planar devices and the simplified single-layer drop generator highlight its ease of operation and robust performance for DE production. This device utilizes core-filling induced drop formation and oil-shearing to reduce the oil volume and ensure the 1:1 core loading of monodisperse DEs without precise synchronization. We probed the stability and coalescence behavior of DE drops during thermal stresses, comparing them with SE drops. DEs resist mixing of contents and eliminate merging, addressing the long-standing challenge of drop stability that has impeded SE-based analyses. Our results show that DEs maintain a monodisperse size distribution by rupturing unstable drops into the continuous phase which effectively removes them. Moreover, DEs prevent drop mixing during merging. This unique coalescence behavior, makes DEs well suited for challenging workflows that have previously been impractical due to SE instability. We demonstrated successful PCR barcoding of viral genomes in drops. Using DEs compartmentalization was preserved, with virtually no barcode mixing, opening up the potential for PCR-based single genome droplet profiling. The results of this effort will help to progress the adoption of powerful DE-based droplet microfluidic techniques, which facilitate lengthy or multi-step reactions in drops, greatly expanding the scope of drop-based assays.

## 2. Experimental

### Fabrication of Microfluidic Devices

Drop generator devices are fabricated using established soft-lithography techniques.^54,55^ A single layer of photoresist (SU-8 2025, 2050) is patterned onto a 3-inch wafer. Optionally, a Karl Suss MJB-3 contact aligner was used to pattern an additional layer of photoresist for two-layer devices. After development, the patterned wafer molds the channels into PDMS slabs (Sylgard). Each PDMS slab is cut from the wafer and device inlet and outlet ports are added using a biopsy punch. The PDMS slabs are treated on high power for 15s at 700 mTorr in a 30W plasma cleaner (Harrick) and bonded together and placed on a glass slide for support. Bonded devices are baked at 65 °C for 48 hrs to restore the native hydrophobic character of the PDMS surface. The devices are sealed with tape and can be stored indefinitely. Immediately before use the tape covering the device outlet is removed and exposed to plasma for 60s which imparts a temporary hydrophilic surface necessary for DE formation. Plasma treatment times will vary with length and cross-section of the outlet channel but can be repeated reliably once an optimized time is determined for each device.

### Double Emulsion Generation

Solutions are loaded into syringe pumps and connected to the device ports using PTFE tubing. When forming DE droplets, the middle oil phase fills the plasma-treated device. Next, the outer aqueous phase is introduced forming a stream of O/W drops at the second junction. This sequence ensures that the OA phase never contacts the serpentine or first junction which would prolong any transient surface oxidation in this region and destabilize W/O drop formation. For visualization purposes, IA consists of 10% green food dye in PBS. The oil phase consists of either the ionic fluorosurfactant Krytox 157 FSH (Miller-Stephenson) or non-ionic 008-fluorosurfactant (Ran Biotechnologies) in HFE-7500 (3M) and is passed through a .2 μm PTFE syringe filter to exclude debris. OA consists of 2% Tween-20 (Fisher) and 1% Poloxamer 188 (Sigma) in 1X PBS (Invitrogen) and is filtered prior to syringe loading. IA, oil, and OA flow rates can be varied over a wide range. Typical flow rates are 125:250:1500 μL hr^-1^ IA:Oil:OA. Drops are collected into 1.5 mL centrifuge tubes.

### Droplet Digital PCR

SV-40 Virus was diluted in molecular biology grade water. PCR mixes consist of 1X Q5 Reaction Buffer, 0.5 μM each forward and reverse primers, 20% GC enhancer, 200 mM dNTP mix, 0.2% Tween-20, 1 unit Q5 Polymerase (New England Biolabs), dilute SV-40, and 0.3X Sybr-Green (Invitrogen). Thermal cycling was performed as follows: 98 °C for 2 min, 45 cycles: (98 °C 10s, 68 °C 20 s, 72 °C 30s), 72 °C for 2 min. Widefield and fluorescent images were obtained using an Axio Observer 5 (Zeiss) inverted microscope.

### Species Mixing Experiment

Separately, PCR mixes consisting of 1X Q5 Reaction Buffer, 20% GC enhancer, 200 μM dNTP mix, 0.1 μM each of BC rev, SV for, λ for primers, 0.025 μM of BC for, SV rev, λ rev primers, 2×10-6 μM barcodes, and 1 unit Q5 Polymerase were prepared along with either SV-40 or Lambda amplicons. Each PCR mix was encapsulated into ~40 μM drops. Following a passion loading of components, ~10% of drops contain a single unique DNA barcode. Each drop contains a few copies of SV-40 or λ. Prior to PCR cycling, Lambda-containing and SV-40 containing droplets are transferred to the same tube. Thermal cycling was performed as follows: 98 °C for 3 min, 25 cycles amplification: (98 °C 30s, 68 °C 30 s, 72 °C 60s), 30 cycles attachment: (98 °C 30s, 59 °C 30 s, 72 °C 60s), 72 °C for 2 min. Single and DE drops were separately diluted with empty drops to reduce the number of barcode positive drops prior to bulk retrieval using 20% 1H,1H,2H,2H-perfluoro-1-octanol (Fisher Scientific) in HFE. Single and DE PCR products are pooled and loaded onto a 1.5% agarose gel. The barcode attached product is recovered by gel purification and P5/P7 adapter sequences are attached: PCR mix consists of gel-purified products, 1X Q5 Reaction Buffer, 20% GC enhancer, 200 μM dNTP mix, 1 μM each P5 and P7 primers, and 1 unit Q5 Polymerase. 98 °C for 3 min, 20 cycles: (98 °C 30s, 68 °C 30 s, 72 °C 60s), 72 °C for 5 min. Products were PCR purified, and confirmed by Sanger sequencing. Prior to sequencing DNA was quantified using Qubit and a bioanalyzer. Sequencing was performed on an Illumina MiSeq Nano. Sequence analysis was performed using a custom R script: Barcodes were identified and grouped with 0-2 mismatches allowed. Then reads were assigned to SV-40 and Lambda reference genomes. SE and DE barcodes were separated using the experiment index included on the barcode primer.

## 3. Results and Discussion

### Simplified Planar Designs for Facile DE Drop Formation

We designed a single-layer PDMS device capable of DE drop formation using two sequential flow-focusing junctions without requiring mask alignment (**Figure 1a,b**). This fully planar device relies on spatial control over the surface wetting to achieve DE drop formation. The single-layer PDMS device is bonded either to a PDMS coated slide or a featureless slab of PDMS resulting in a channel surface that is entirely PDMS. After device bonding, hydrophobic recovery of the PDMS^53^ ensures that the first junction will properly encapsulate the inner aqueous flow in oil. Immediately prior to use, a simple plasma treatment applies a transient hydrophilic surface to the second junction and device outlet. To achieve spatial patterning, device inlets are covered with tape and plasma treatment times are minimized such that plasma diffuses only into the outlet and travels towards the junction applying the hydrophilic surface wetting. **Figure 1c** provides an example of the DEs produced using this device. The resulting DEs are highly monodisperse with the coefficient of variation <5%. By eliminating unnecessary complexity, this alignment-free device requires only a single layer PDMS device and a straightforward surface treatment to generate monodisperse DE drops, reducing the barriers to the adoption of DE techniques.

**Figure 1.**
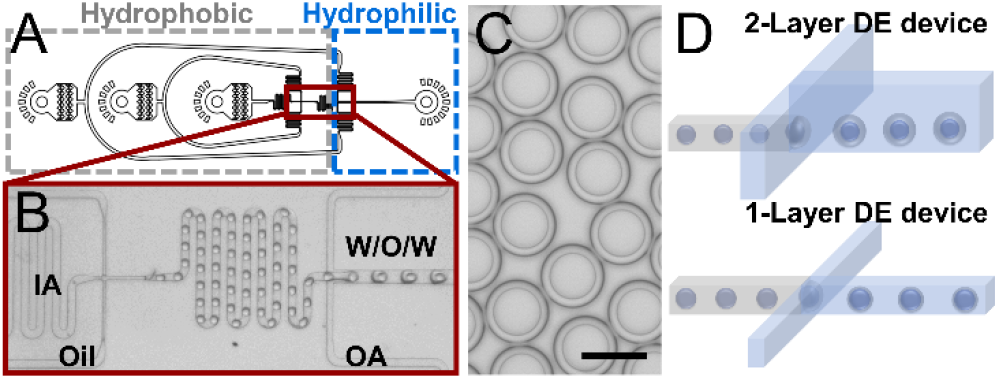
A) Device design and surface wetting of the single-layer DE drop maker. B) The inner aqueous phase (IA) is encapsulated by the oil to form a water-in-oil (W/O) emulsion at the hydrophobic junction. Subsequently, the outer aqueous phase (OA) breaks up this stream into a water-in-oil-in-water (W/O/W) DE at the hydrophilic junction. C) Monodisperse DEs generated using a single layer device. The scale bar is 50 μm. D) Schematic representation of a standard two-layer DE device with a height change at the second junction and the simplified one-layer DE device.

### Robust Double Emulsion Generation in a Fully Planar Device

To compare the performance of the newly designed single-layer device, a conventional semi-planar DE drop maker was fabricated featuring a height increase at the second junction (**Figure 1d**). Surprisingly, our simplified planar device was able to robustly produce monodisperse DE drops over a wide range of flow rates, while the semi-planar device required precise flow rate optimization to generate high-quality DEs (**Figure 2**). In both cases, the inner aqueous (IA) and oil flow rate were stabilized to form a stream of water-in-oil (W/O) drops. The outer aqueous phase (OA) flow rate was varied and double emulsification was assessed. For both devices, the OA flow rate must be above a critical threshold, to break up the stream of SE drops into DEs. However, for the 2-layer device at low OA flow rates, pinch off of the filling DE drop occurs slowly and multiple aqueous cores are being loaded into the filling DE drop before it pinches off (**Figure 2a**, 450 μL hr^-1^). As the outer flow rate is increased, the proportion of DE drops with two cores decreases. For the two-layer device, a narrow range of flow rates results in optimized aqueous core loading to form 1:1 DEs (**Figure 2a**, 750 μL hr^-1^). As the OA flow rate is increased beyond this optimized flow regime, drop pinch-off can occur as an aqueous core is entering the filling DE drop, resulting in partial core loading, which is not suitable for most applications (**Figure 2a**, 950 μL hr^-1^). Conversely, for the single-layer device, single-core DEs are formed over a wide range of flow rates spanning nearly an order of magnitude (**Figure 2d**) before partial core loading begins. The fully planar geometry of the device facilitates this unique stability in drop formation behavior. Because the channel height and width remain constant at the second junction, the arrival of each aqueous core to the filling DE drop induces pinch-off. This occurs because the aqueous cores contain a substantial fraction of volume required for drop formation in the dripping regime,^56^ so the instability of the growing DE cannot be sustained upon core filling inducing drop formation in this confined channel geometry. This allows for 1:1 DE drop loading without synchronization of sequential drop generation steps. This significantly reduces the difficulty of DE drop generation and removes most laborious flow rate optimizations. **Figure 2e** shows the broad range of suitable flow rates for high-quality DE generation. These results illustrate the ease of device operation afforded by the single-layer design. In both cases, the aqueous core and shell diameters are highly monodisperse within the optimized DE formation regime, but the single-layer device is able to maintain excellent monodispersity and single-core encapsulation across a broad range of flow conditions. Using this planar design to simplify device fabrication and operation, DE drops generation is now comparable to SE generation, which sees routine use.

**Figure 2.**
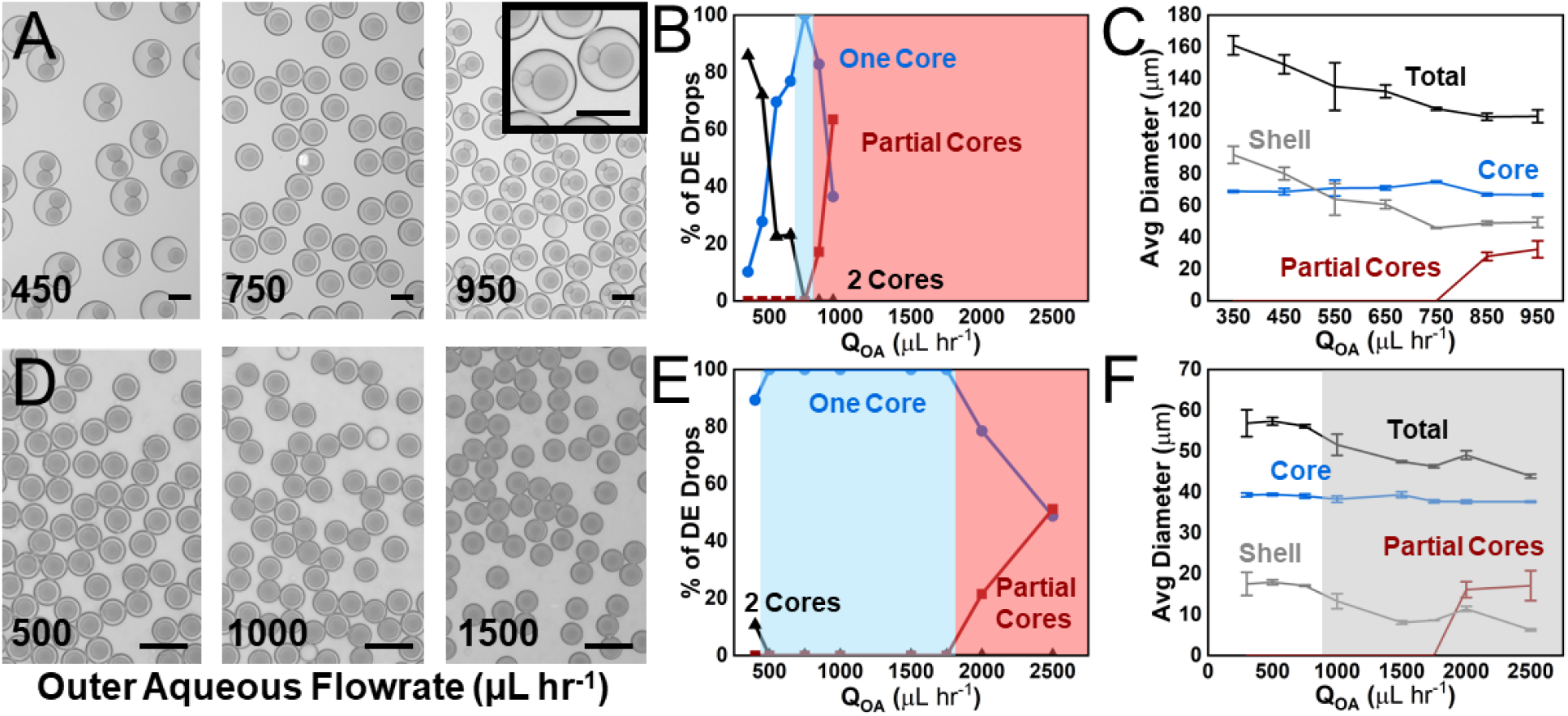
A) Representative images of DE drops formed at 450, 750, 950 μL hr^-1^. The scale bar is 100 μm. B) Flow rate dependence of core loading of aqueous drops into DEs. Using a standard 2 layer-device geometry, DE generation is highly sensitive to the chosen flow rates, and 1:1 DE drop formation occurs only within a narrow range. C) Flow rate dependence of DE sizes. Shell diameter is defined as the difference between the total DE drop and core diameter. D) Using a single-layer device design DE generation occurs robustly across a wide range of flow rates. Microscope images of resulting DEs are shown at 500, 1000, 1500 μL hr^-1^. The scale bar is 100 μm. E) Flow rate dependence of core loading of aqueous drops into DEs. The confined channel at the second junction ensures DE drops are loaded 1:1. F) Flow rate dependence of core and shell diameters. Shell thicknesses significantly reduced when compared with DE drops produced from a 2-layer device. Further reduction in shell-thickness occurs due to on-chip shearing in the shaded region.

### On-chip Shearing Controls Shell Monodispersity

The confined channels of the planar design yield a reduction in oil shell thickness compared to the 2-layer semi-planar device. For both designs, increases in the OA flow rate resulted in thinner shells (**Figure 2c,f**). The single-layer device, however, produces DEs with thinner shells relative to the core diameter. Thin shells are desirable for FACS sorting of DE drops, as excess oil volumes can shear within the instrument and cause clogging. Reduced shell volumes improve FACS sorting and have use for microcapsule fabrication.^57–60^ At increased OA flow rates, the single-layer device design drives thin shell formation *via* a distinct mechanism. Using a high-speed camera, we observed on-chip shearing of excess oil as the formed DE drops traveled down the confined channel towards the outlet (**Figure 3a**). Above a critical flow rate, this oil shearing stabilizes such that all DE shells are sheared equally resulting in the uniform generation of monodisperse DEs with thin shells (**Figure 2f, S2**). In this flow regime, core arrivals are not synchronized with the oil filling of the DE drop. Accordingly, DE drops can pinch off with different oil volumes but are sheared uniformly in the flow (**Supplementary Video 1**). This mechanism enables monodisperse shell thicknesses despite a lack of synchronized drop formation. Semi-planar DE drop generation designs include a channel height increase at the second junction which prevents both core-filling induced drop formation and on-chip oil shearing. The DE generation comparisons between 1- and 2-layer devices emphasize the ease of operation enabled by the planar DE drop maker and its robust ability to produce monodisperse DE drops without synchronizing the drop generation at the first and the second junctions.

**Figure 3.**
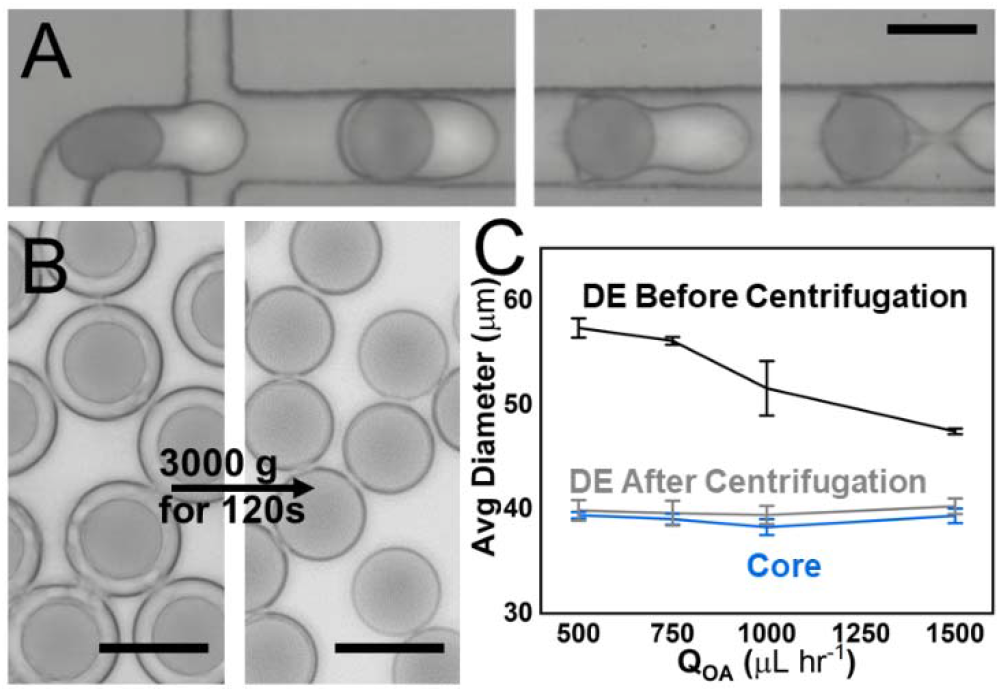
A) Sequential images of a DE drop during on-chip shearing. B) Microscope images showing ultra-thin shelled DE production by centrifugation. Scale bars are 50 μm. C) Original DEs were produced at flow rates of IA:Oil:OA=125:250:[500, 1000, 1500, 1750] μL hr^-1^. Following centrifugation-induced shearing, the resulting DEs are monodisperse.

### Centrifugation of Double Emulsions for Oil Separation

On-chip shearing results in the formation of excess oil droplets, which could potentially impact certain downstream applications like drop-by-drop screening, by preventing uniform close-packing of the emulsion. However, SE oil drops and DEs have significantly different densities, so they can be separated using a bench-top centrifuge. The DEs are easily recovered by pipette and transferred to a clean tube with excess continuous phase. Centrifugation at low speeds results in effective separation with minimal drop rupture and no change in the oil shell thickness (**Figure S3**). This simple centrifugation method reliably separates DE drops from excess oil drops, ensuring that the advantages of these simplified DE formation methods are not impeded by additional oil drops.

We further hypothesized that higher centrifugation speeds would induce shearing of the oil shells, and eventually droplet coalescence. Previous efforts to make DEs with extremely thin shells have required complex microfluidic structures or flow optimization to controllably extract oil from DEs.^61,62^ To drive further shearing of the oil shell while minimizing DE rupture, we optimized the centrifugation speed. As expected, DE drops with ultra-thin shells were readily produced using this approach (**Figure 3b**). After centrifugation at 3000g for 2 minutes, the oil shell thickness is reduced significantly. The post-centrifugation shell thickness was independent of the starting shell thickness (**Figure 3c**) leaving behind only a minimal amount of oil around each aqueous core. Because the shell thickness was so small it is no longer feasible to measure optically. Instead, isopropanol was added to the DE drops inducing rupture (**Figure S4a**). The fluorinated oil that makes up the shell coalesces into a single oil drop upon rupture, which is of sufficient size to be measured. Approximately 50 ultrathin shelled DEs were assessed in this way (**Figure S4b**). Following centrifugation and shearing, DE drops had an average diameter of 41.1 ± 1.5 μm. Upon rupture in IPA, shell content drops were 12.5 ± 1.6 μm in diameter. Assuming two concentric spheres, this gives an average shell thickness of 193 nm. This method illustrates the potential to obtain DEs with ultra-thin shells in a simple and high-throughput way using a common benchtop centrifuge. This method further expands the potential applications for DE drops.

### Double Emulsions Show Enhanced Stability During Droplet Digital PCR

DEs show potential as an alternative to SEs for droplet PCR due to their impressive stability. Droplet digital PCR,^63^ in-drop directed evolution,^64^ and other applications routinely utilize PCR amplification in emulsion drops. Thermal cycling procedures, however, apply stress to emulsions, which inevitably induce coalescence. Considerable effort has been dedicated to designing and optimizing surfactants capable of stabilizing emulsions during PCR but drop merging still remains a frustrating challenge for SE microfluidics.^7,8^ Even specialized surfactants cannot prevent merging entirely, and often the extent of merging can vary in ways that are difficult to predict. Often, the degree of drop merging becomes worse as surfactants age and significant batch-to-batch variation exists for commercially available surfactants. **Figure S1** shows the widely varying extent of merging that results from three replicate ddPCR experiments. Merging not only forms larger drops, but also a completely coalesced layer of aqueous material as seen in **Figure S1b,c**. Material from this merged layer would interfere with drop analysis. The differences in drop coalescence between SEs and DEs have not previously been studied in detail, but our results indicate that DE drops are more robust during PCR thermal cycling. Larger drops are less stable, so SE PCR approaches typically use smaller drop sizes (< 50 μm) to reduce merging. However, smaller drops also limit the maximum total yield of each reaction in drops. To test the potential for DE drops to facilitate PCR amplification or other reaction steps in larger drops, we generated ~80 μm SEs and DEs and performed ddPCR. Each emulsion drop consists of an aqueous volume of a PCR mixture containing dilute SV-40 virus and SYBR green dye. Since the viral genomes are sufficiently diluted each droplet contains either 0 or 1 copies resulting in digital amplification during PCR. The droplets that contain a DNA template amplify during PCR and fluoresce in the presence of SYBR-green dyes. **Figure 4** shows the results large drop ddPCR in SEs and DEs.

**Figure 4.**
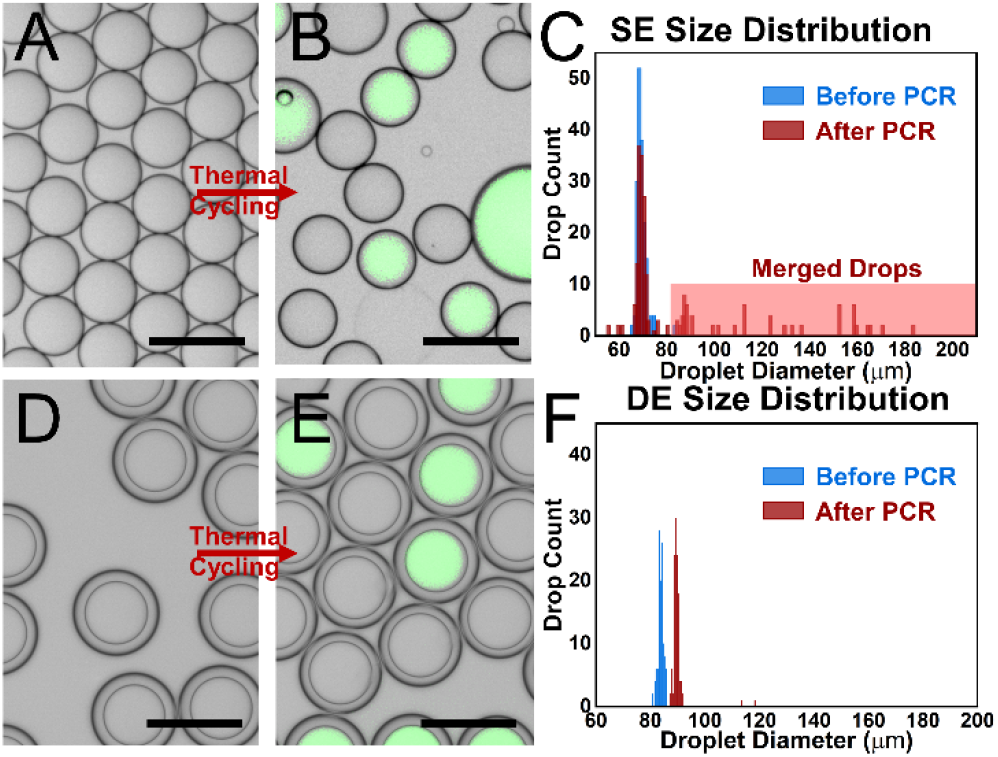
Comparison between ~80 μm SEs and DEs during ddPCR. A, D) Drops after collection. B, E) Drops after thermal cycling. SYBR-green fluorescence indicates ddPCR amplification. The scale bar is 100 μm. C, F) Histograms summarizing drop size distributions throughout the ddPCR workflow. The DE maintains a monodisperse size distribution while SE drops merge to form larger drops.

In both cases, single virus genomes are successfully amplified in drops. Notably, the SE drops undergo substantial merge during PCR resulting in the formation of larger drops (**Figure 4b,c**) and a merged aqueous layer (**Figure S5a**). In contrast, the DE maintains a monodisperse size distribution after thermal cycling (**Figure 4e,f**) and mostly survives thermal cycling (**Figure S5b**). Monodisperse drops are ideal for quantification by ddPCR as the proportion of bright drops can be easily related to a concentration. Although polydisperse emulsions can be analyzed for ddPCR quantification, the measurement error is greater.^65^ This experiment was repeated with ~40 μm SE and DE drops. As expected, the total amount of merging was less at this smaller drop size, but the SE still displayed some merging (**Figure S6**). The DE showed no change in the drop distribution following PCR. This improvement in emulsion stability enables DEs to perform reactions in larger droplets or to withstand repeated thermal stresses.

### Compartmentalization of Contents

Beyond the impacts to the size distribution, drop merging also has the potential to undesirably mix drop contents during PCR amplification or other reactions in drops. Droplet emulsions are routinely used for single-cell analysis, which necessitates isolation of drop contents for error-free analysis. To investigate the effect of droplet merging on drop contents, SEs and DEs were prepared and labeled using two dyes. This allows for the straightforward visualization of the compartmentalization of drop contents. The difference in coalescence behavior between SEs and DEs can be seen in **Figure 5**. For SEs, drop merging fuses aqueous volumes forming larger drops. Critically, merging generates big drops that contain a mix of both dye colors following PCR indicating a breakdown of compartmentalization. DEs respond differently upon coalescence, rupturing their aqueous contents into the continuous aqueous phase where no reagents are present. Outer aqueous phase material is removed from downstream analysis. Accordingly, DEs are able to fully compartmentalize each dye population and prevent cores from mixing even when drop coalescence occurs. These effects were also observed in heterogeneous drop populations (**Figure S7a,b**). Again, the DE drops effectively maintained compartmentalization.

**Figure 5.**
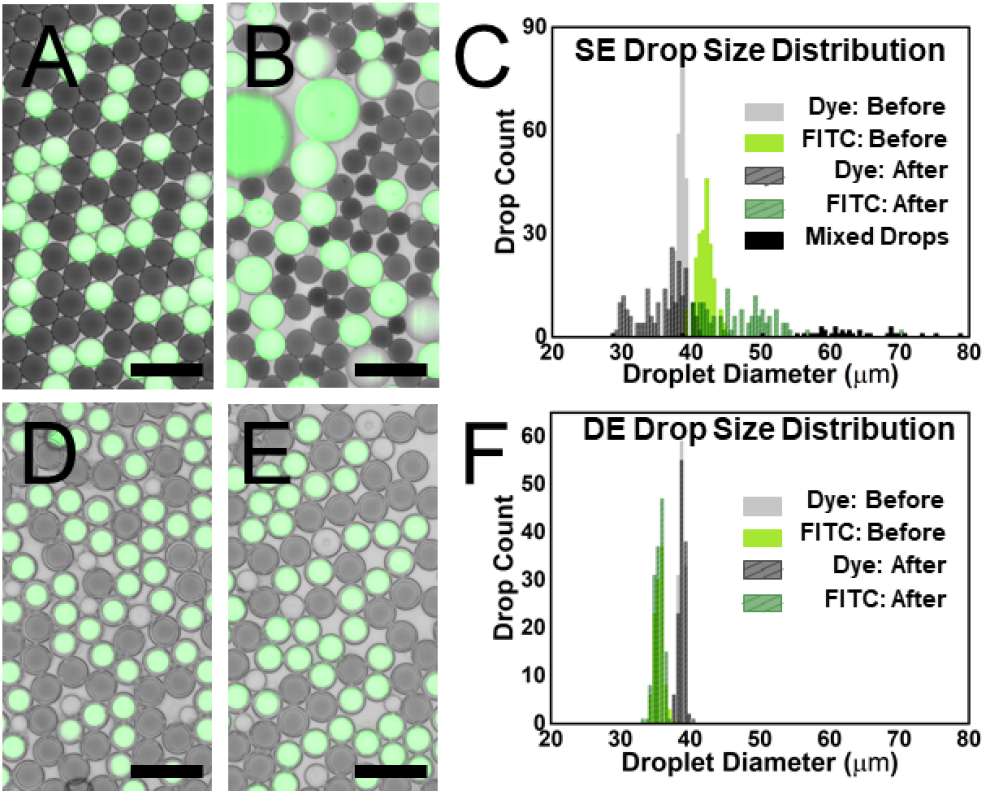
Comparison between PCR merging behavior of SEs and DEs. A, D) Microscope images of mixed ~40 μm drops containing different dyes. B, E) Images of drops after thermal cycling. The scale bar is 100 μm. C, F) Histograms summarizing drop size distribution before and after PCR induced merging. DEs maintain a monodisperse size distribution and prevent mixing of their aqueous contents while SE drops merge and mix their contents.

This coalescence behavior is a key strength of DEs for single entity analysis. To illustrate this, we performed targeted PCR barcoding in drops as a simplified version of a PCR-based single genome screening assay. In this scheme, unique DNA barcodes are loaded into droplets containing one of two viral targets where they attach to a portion of the genome and amplify. Each DNA barcode has a unique sequence that represents the droplet in which it is loaded. During PCR, barcodes attach to the amplicons present in the drop providing a record of the sequences that we compartmentalized. Specifically, amplification and barcoding were achieved using a set of primers designed to target specific regions within either the SV-40 or λ-phage genome. A shared handle sequence on each primer allows the drop-specific barcodes to attach to either amplicon during PCR. This mimics any protocol that uses PCR to label genomic or transcriptomic elements originating from single cells inside drops. Drop populations were prepared separately to ensure that each droplet contains only one of the two virus types. The two drop populations were then combined into the same tube prior to thermal cycling. In this way, we ensure that the input drops compartmentalize either SV-40 or λ-phage sequences but not both. During PCR, drop coalescence occurs which has the potential to disrupt this compartmentalization. Sequencing libraries were prepared using either SE or DE drops and the results were compared by analyzing the distribution of reads mapping to the drop-specific barcodes. If compartmentalization is maintained, each barcode should be associated with reads from only SV-40 or λ but not both. As expected, SE drops undergo merging which results in droplet-specific barcodes with reads from both targets (**Figure 6c**), which is detrimental for single genome analysis. In contrast, while the DEs effectively prevent mixing even as some drops are ruptured (**Figure 6d**). Each DE barcode displays reads from either SV-40 or λ-phage but not both. This confirms that DE drops are able to effectively maintain compartmentalization even as some drops inevitably rupture. Accordingly, DEs provide a crucial modality to develop PCR-based assays targeting single cells or viruses in drops. Unlike SEs, the contents in DE drops that coalesce are simply expelled into the inert outer aqueous phase effectively removing ruptured drops from downstream analysis. The ruptured contents can be readily removed from the system by transferring intact emulsions to new buffers prior to bulk recovery or further manipulation of drops. The ability of DEs to avoid content mixing ensures that compartmentalization is robust especially during lengthy and high-stress procedures, where this enhanced stability is a crucial advantage.

**Figure 6.**
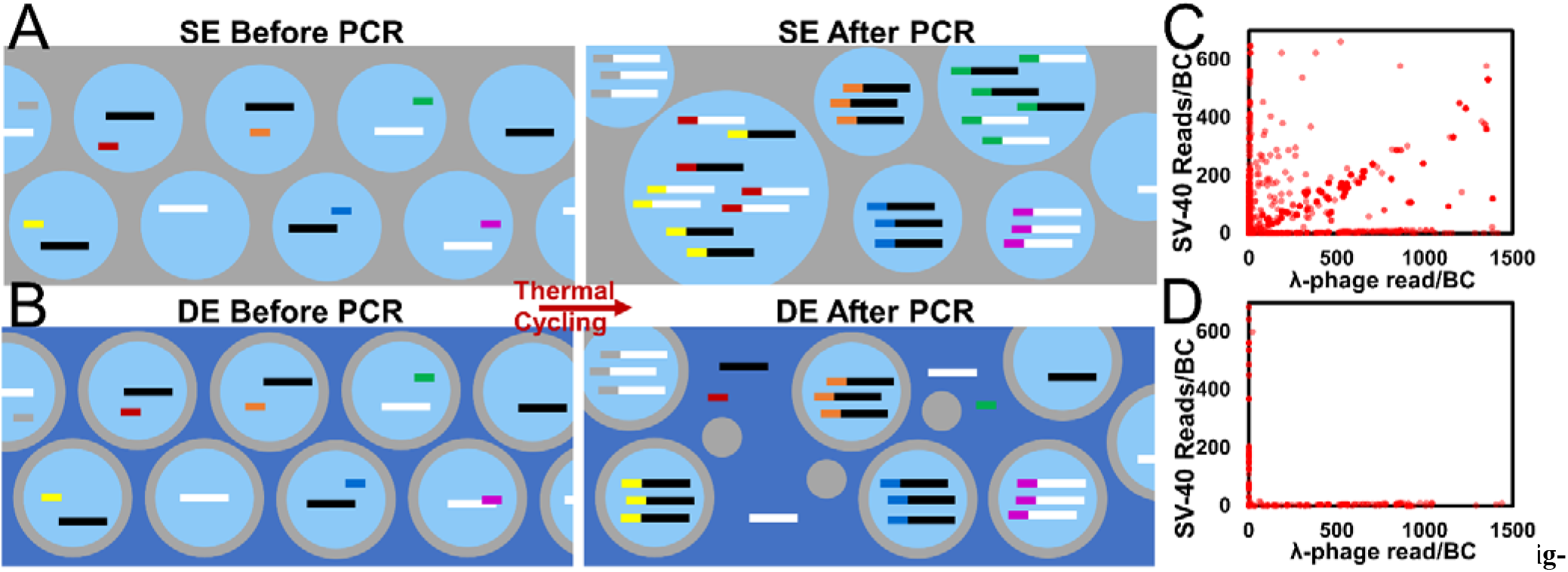
A) Schematic representation of PCR barcoding experiment. Unique drop-specific barcodes amplify and attach to all amplicons present during PCR. Plot of SV-40 and λ-Phage reads per SE (C) or DE (D) drop barcode. Some SE barcodes are observed with reads from both species. Each DE barcode has only reads from one virus type indicating effective compartmentalization.

## 4. Conclusions

These results show the utility of DEs as a robust alternative to SE drops. Using simplified single-layer devices and surface treatments, it is possible to generate high-quality monodisperse DEs without precisely optimizing flow conditions. Using core-filling-induced drop formation and on-chip shearing, monodisperse DE drops with desirable thin shells are formed over a broad range of flow conditions. Using this simplified design, it is now much easier to utilize the increased stability and robust compartmentalization afforded by DE drops. This work lays the foundation for the broader use of DE drops for PCR-based methods, flow cytometry sorting, and more. Coalescence comparisons investigated the stability and merging dynamics of SE and DE drops. Our results indicate that DEs coalesce primarily through drop rupture. Unlike merging in SEs, drop rupture ensures that DE drop contents do not mix and the emulsion remains monodisperse even as some portion coalesces. In applications where drop mixing must be fully eliminated, DEs provide a clear solution, ensuring that merging does not contaminate downstream processing and analysis. Moreover, DE drops retain their monodispersity even across lengthy and multistep processes and at larger drop sizes. Maintaining a uniform drop size is important as it ensures that each reaction vessel is loaded with similar reagent volumes improving the uniformity of yields across drops. This is particularly advantageous for sequencing experiments, where excessive amplification in merged drops will waste reads. Moreover, drop monodispersity is a key requirement for deterministic drop-by-drop manipulations. This study not only demonstrates the unique strength of DE drops to withstand coalescence but also introduces a simple and robust method to produce DE drops as easily as SE drops. As shown, DEs have the potential to expand the scope of routine microfluidic assays to incorporate more demanding steps that SE drops are unable to withstand.

## ASSOCIATED CONTENT

### Supporting Information

The Supporting Information is available free of charge on the publications website.

Supporting information (pdf)

Video of On-Chip Shearing (mov)

## Author Contributions

T.W.C., and H.-S.H. have conceived of the idea and designed experiments. T.W.C. designed drop generation devices. T.W.C. and A.D. fabricated drop generation devices, made drops, performed PCR, collected images, and analyzed data. T.W.C. and H.-S.H prepared the manuscript.

## Notes

The authors declare no other competing financial interests.

## ACKNOWLEDGMENTS

This work was carried out in part in the Materials Research Laboratory Central Research Facilities, University of Illinois. Sequencing was performed by the Roy J. Carver Biotechnology Center. The authors also thank the University of Illinois at Urbana-Champaign and the Gordon and Betty Moore Foundation for their support.

## REFERENCES

(1) Teh, S. Y.; Lin, R.; Hung, L. H.; Lee, A. P. Droplet Microfluidics. Lab on a Chip. The Royal Society of Chemistry January 29, 2008, pp 198–220. https://doi.org/10.1039/b715524g.

(2) Mazutis, L.; Gilbert, J.; Ung, W. L.; Weitz, D. A.; Griffiths, A. D.; Heyman, J. A. Single-Cell Analysis and Sorting Using Droplet-Based Microfluidics. Nat. Protoc. 2013, 8 (5), 870–891. https://doi.org/10.1038/nprot.2013.046.

(3) Klein, A. M.; Mazutis, L.; Akartuna, I.; Tallapragada, N.; Veres, A.; Li, V.; Peshkin, L.; Weitz, D. A.; Kirschner, M. W. Droplet Barcoding for Single-Cell Transcriptomics Applied to Embryonic Stem Cells. Cell 2015, 161 (5), 1187–1201. https://doi.org/10.1016/j.cell.2015.04.044.

(4) Stucki, A.; Vallapurackal, J.; Ward, T. R.; Dittrich, P.S. Droplet Microfluidics and Directed Evolution of Enzymes: An Intertwined Journey. Angew. Chemie Int. Ed. 2021. https://doi.org/10.1002/anie.202016154.

(5) Macosko, E. Z.; Basu, A.; Satija, R.; Nemesh, J.; Shekhar, K.; Goldman, M.; Tirosh, I.; Bialas, A. R.; Kamitaki, N.; Martersteck, E. M.; et al. Highly Parallel Genome-Wide Expression Profiling of Individual Cells Using Nanoliter Droplets. Cell 2015, 161 (5), 1202–1214. https://doi.org/10.1016/j.cell.2015.05.002.

(6) Chiu, F. W. Y.; Stavrakis, S. High-Throughput Droplet-Based Microfluidics for Directed Evolution of Enzymes. Electrophoresis. John Wiley & Sons, Ltd November 1, 2019, pp 2860–2872. https://doi.org/10.1002/elps.201900222.

(7) Payne, E. M.; Holland-Moritz, D. A.; Sun, S.; Kennedy, R. T. High-Throughput Screening by Droplet Microfluidics: Perspective into Key Challenges and Future Prospects. Lab on a Chip. 2020, pp 2247–2262. https://doi.org/10.1039/d0lc00347f.

(8) Chowdhury, M. S.; Zheng, W.; Kumari, S.; Heyman, J.; Zhang, X.; Dey, P.; Weitz, D. A.; Haag, R. Dendronized Fluorosurfactant for Highly Stable Water-in-Fluorinated Oil Emulsions with Minimal Inter-Droplet Transfer of Small Molecules. Nat. Commun. 2019, 10 (1), 1–10. https://doi.org/10.1038/s41467-019-12462-5.

(9) Holtze, C.; Rowat, A. C.; Agresti, J. J.; Hutchison, J. B.; Angilè, F. E.; Schmitz, C. H. J.; Köster, S.; Duan, H.; Humphry, K. J.; Scanga, R. A.; et al. Biocompatible Surfactants for Water-in-Fluorocarbon Emulsions. Lab Chip 2008, 8 (10), 1632–1639. https://doi.org/10.1039/b806706f.

(10) Cheng, G.; Lin, K. T.; Ye, Y.; Jiang, H.; Ngai, T.; Ho, Y. P. Photo-Responsive Fluorosurfactant Enabled by Plasmonic Nanoparticles for Light-Driven Droplet Manipulation. ACS Appl. Mater. Interfaces 2021, 13 (18), 21914–21923. https://doi.org/10.1021/acsami.0c22900.

(11) Goodarzi, F.; Zendehboudi, S. A Comprehensive Review on Emulsions and Emulsion Stability in Chemical and Energy Industries. Can. J. Chem. Eng. 2019, 97 (1), 281–309. https://doi.org/10.1002/cjce.23336.

(12) Ooi, Z. Y.; Othman, N.; Choo, C. L. The Role of Internal Droplet Size on Emulsion Stability and the Extraction Performance of Kraft Lignin Removal from Pulping Wastewater in Emulsion Liquid Membrane Process. J. Dispers. Sci. Technol. 2016, 37 (4), 544–554. https://doi.org/10.1080/01932691.2015.1050728.

(13) Oren, J. J.; MacKay, G. D. M. Electrolyte and PH Effect on Emulsion Stability of Water-in-Petroleum Oils. Fuel 1977, 56 (4), 382–384. https://doi.org/10.1016/0016-2361(77)90062-X.

(14) Zeeb, B.; Zhang, H.; Gibis, M.; Fischer, L.; Weiss, J. Influence of Buffer on the Preparation of Multilayered Oil-in-Water Emulsions Stabilized by Proteins and Polysaccharides. Food Res. Int. 2013, 53 (1), 325–333. https://doi.org/10.1016/j.foodres.2013.04.017.

(15) Dogan, M.; Göksel Saraç, M.; Aslan Türker, D. Effect of Salt on the Inter-Relationship between the Morphological, Emulsifying and Interfacial Rheological Properties of O/W Emulsions at Oil/Water Interface. J. Food Eng. 2020, 275, 109871. https://doi.org/10.1016/j.jfoodeng.2019.109871.

(16) Binks, B. P.; Rocher, A. Effects of Temperature on Water-in-Oil Emulsions Stabilised Solely by Wax Microparticles. J. Colloid Interface Sci. 2009, 335 (1), 94–104. https://doi.org/10.1016/j.jcis.2009.03.089.

(17) Perumanath, S.; Borg, M. K.; Chubynsky, M. V.; Sprittles, J. E.; Reese, J. M. Droplet Coalescence Is Initiated by Thermal Motion. Phys. Rev. Lett. 2019, 122 (10), 104501. https://doi.org/10.1103/PhysRevLett.122.104501.

(18) Luhede, L.; Wollborn, T.; Fritsching, U. Stability of Multiple Emulsions under Shear Stress. Can. J. Chem. Eng. 2020, 98 (1), 186–193. https://doi.org/10.1002/cjce.23578.

(19) Abate, A. R.; Hung, T.; Marya, P.; Agresti, J. J.; Weitz, D. A. High-Throughput Injection with Microfluidics Using Picoinjectors Using Picoinjectors. Proc. Natl. Acad. Sci. U. S. A. 2010, 107 (45), 19163–19166. https://doi.org/10.1073/pnas.1006888107.

(20) Zheng, G. X. Y.; Terry, J. M.; Belgrader, P.; Ryvkin, P.; Bent, Z. W.; Wilson, R.; Ziraldo, S. B.; Wheeler, T. D.; McDermott, G. P.; Zhu, J.; et al. Massively Parallel Digital Transcriptional Profiling of Single Cells. Nat. Commun. 2017, 8 (1), 1–12. https://doi.org/10.1038/ncomms14049.

(21) Hindson, B. J.; Ness, K. D.; Masquelier, D. A.; Belgrader, P.; Heredia, N. J.; Makarewicz, A. J.; Bright, I. J.; Lucero, M. Y.; Hiddessen, A. L.; Legler, T. C.; et al. High-Throughput Droplet Digital PCR System for Absolute Quantitation of DNA Copy Number. Anal. Chem. 2011, 83 (22), 8604–8610. https://doi.org/10.1021/ac202028g.

(22) Sukovich, D. J.; Lance, S. T.; Abate, A. R. Sequence Specific Sorting of DNA Molecules with FACS Using 3dPCR. Sci. Rep. 2017, 7 (1), 1–9. https://doi.org/10.1038/srep39385.

(23) Brower, K. K.; Carswell-Crumpton, C.; Klemm, S.; Cruz, B.; Kim, G.; Calhoun, S. G. K.; Nichols, L.; Fordyce, P. M. Double Emulsion Flow Cytometry with High-Throughput Single Droplet Isolation and Nucleic Acid Recovery. Lab Chip 2020, 20 (12), 2062–2074. https://doi.org/10.1039/D0LC00261E.

(24) Brower, K. K.; Khariton, M.; Suzuki, P. H.; Still, C.; Kim, G.; Calhoun, S. G. K.; Qi, L. S.; Wang, B.; Fordyce, P. M. Double Emulsion Picoreactors for High-Throughput Single-Cell Encapsulation and Phenotyping via FACS. Anal. Chem. 2020, 92 (19), 13262–13270. https://doi.org/10.1021/acs.analchem.0c02499.

(25) Mazutis, L.; Gilbert, J.; Ung, W. L.; Weitz, D. A.; Griffiths, A. D.; Heyman, J. A. Single-Cell Analysis and Sorting Using Droplet-Based Microfluidics. Nat. Protoc. 2013, 8 (5), 870–891. https://doi.org/10.1038/nprot.2013.046.

(26) Vladisavljević, G. T.; Al Nuumani, R.; Nabavi, S. A. Microfluidic Production of Multiple Emulsions. Micromachines. Multidisciplinary Digital Publishing Institute (MDPI) 2017. https://doi.org/10.3390/mi8030075.

(27) Shah, R. K.; Shum, H. C.; Rowat, A. C.; Lee, D.; Agresti, J. J.; Utada, A. S.; Chu, L. Y.; Kim, J. W.; Fernandez-Nieves, A.; Martinez, C. J.; et al. Designer Emulsions Using Microfluidics. Materials Today. Elsevier April 1, 2008, pp 18–27. https://doi.org/10.1016/S1369-7021(08)70053-1.

(28) Chong, D. T.; Liu, X. S.; Ma, H. J.; Huang, G. Y.; Han, Y. L.; Cui, X. Y.; Yan, J. J.; Xu, F. Advances in Fabricating Double-Emulsion Droplets and Their Biomedical Applications. Microfluidics and Nanofluidics. Springer September 9, 2015, pp 1071–1090. https://doi.org/10.1007/s10404-015-1635-8.

(29) Nabavi, S. A.; Vladisavljević, G. T.; Gu, S.; Ekanem, E. E. Double Emulsion Production in Glass Capillary Microfluidic Device: Parametric Investigation of Droplet Generation Behaviour. Chem. Eng. Sci. 2015, 130, 183–196. https://doi.org/10.1016/j.ces.2015.03.004.

(30) Sattari, A.; Hanafizadeh, P. Controllable Preparation of Double Emulsion Droplets in a Dual-Coaxial Microfluidic Device. J. Flow Chem. 2021, 1–15. https://doi.org/10.1007/s41981-021-00155-4.

(31) Michelon, M.; Huang, Y.; de la Torre, L. G.; Weitz, D. A.; Cunha, R. L. Single-Step Microfluidic Production of W/O/W Double Emulsions as Templates for β-Carotene-Loaded Giant Liposomes Formation. Chem. Eng. J. 2019, 366, 27–32. https://doi.org/10.1016/j.cej.2019.02.021.

(32) Hou, L.; Ren, Y.; Jia, Y.; Deng, X.; Tang, Z.; Tao, Y.; Jiang, H. A Simple Microfluidic Method for One-Step Encapsulation of Reagents with Varying Concentrations in Double Emulsion Drops for Nanoliter-Scale Reactions and Analyses. Anal. Methods 2017, 9 (17), 2511–2516. https://doi.org/10.1039/c7ay00544j.

(33) Martino, C.; Berger, S.; Wootton, R. C. R.; Demello, A. J. A 3D-Printed Microcapillary Assembly for Facile Double Emulsion Generation. Lab Chip 2014, 14 (21), 4178–4182. https://doi.org/10.1039/c4lc00992d.

(34) Beauchamp, M. J.; Gong, H.; Woolley, A. T.; Nordin, G. P. 3D Printed Microfluidic Features Using Dose Control in X, Y, and Z Dimensions. Micromachines 2018, 9 (7), 326. https://doi.org/10.3390/mi9070326.

(35) Sanchez Noriega, J. L.; Chartrand, N. A.; Valdoz, J. C.; Cribbs, C. G.; Jacobs, D. A.; Poulson, D.; Viglione, M. S.; Woolley, A. T.; Van Ry, P. M.; Christensen, K. A.; et al. Spatially and Optically Tailored 3D Printing for Highly Miniaturized and Integrated Microfluidics. Nat. Commun. 2021, 12 (1), 1–13. https://doi.org/10.1038/s41467-021-25788-w.

(36) Romanowsky, M. B.; Abate, A. R.; Rotem, A.; Holtze, C.; Weitz, D. A. High Throughput Production of Single Core Double Emulsions in a Parallelized Microfluidic Device. Lab Chip 2012, 12 (4), 802–807. https://doi.org/10.1039/c2lc21033a.

(37) Lim, S. W.; Abate, A. R. Ultrahigh-Throughput Sorting of Microfluidic Drops with Flow Cytometry. Lab Chip 2013, 13 (23), 4563–4572. https://doi.org/10.1039/c3lc50736j.

(38) Alsharhan, A. T.; Stair, A. J.; Acevedo, R.; Razaulla, T.; Warren, R.; Sochol, R. D. Direct Laser Writing for Deterministic Lateral Displacement of Submicron Particles. J. Microelectromechanical Syst. 2020, 29 (5), 906–911. https://doi.org/10.1109/JMEMS.2020.2998958.

(39) Niculescu, A. G.; Chircov, C.; Bîrcă, A. C.; Grumezescu, A. M. Fabrication and Applications of Microfluidic Devices: A Review. International Journal of Molecular Sciences. Multidisciplinary Digital Publishing Institute February 18, 2021, pp 1–26. https://doi.org/10.3390/ijms22042011.

(40) Scott, S. M.; Ali, Z. Fabrication Methods for Microfluidic Devices: An Overview. Micromachines. Multidisciplinary Digital Publishing Institute (MDPI) March 1, 2021. https://doi.org/10.3390/mi12030319.

(41) Lee, T. Y.; Choi, T. M.; Shim, T. S.; Frijns, R. A. M.; Kim, S. H. Microfluidic Production of Multiple Emulsions and Functional Microcapsules. Lab on a Chip. The Royal Society of Chemistry August 30, 2016, pp 3415–3440. https://doi.org/10.1039/c6lc00809g.

(42) Ma, S.; Huck, W. T. S.; Balabani, S. Deformation of Double Emulsions under Conditions of Flow Cytometry Hydrodynamic Focusing. Lab Chip 2015, 15 (22), 4291–4301. https://doi.org/10.1039/c5lc00693g.

(43) Nabavi, S. A.; Vladisavljević, G. T.; Manović, V. Mechanisms and Control of Single-Step Microfluidic Generation of Multi-Core Double Emulsion Droplets. Chem. Eng. J. 2017, 322, 140–148. https://doi.org/10.1016/J.CEJ.2017.04.008.

(44) Zhang, J. Q.; Chang, K.-C.; Liu, L.; Gartner, Z. J.; Abate, A. R. High Throughput Yeast Strain Phenotyping with Droplet-Based RNA Sequencing. J. Vis. Exp. 2020, No. 159, e61014. https://doi.org/10.3791/61014.

(45) Habib, N.; Avraham-Davidi, I.; Basu, A.; Burks, T.; Shekhar, K.; Hofree, M.; Choudhury, S. R.; Aguet, F.; Gelfand, E.; Ardlie, K.; et al. Massively Parallel Single-Nucleus RNA-Seq with DroNc-Seq. Nat. Methods 2017, 14 (10), 955–958. https://doi.org/10.1038/nmeth.4407.

(46) Feng, Y.; White, A. K.; Hein, J. B.; Appel, E. A.; Fordyce, P. M. MRBLES 2.0: High-Throughput Generation of Chemically Functionalized Spectrally and Magnetically Encoded Hydrogel Beads Using a Simple Single-Layer Microfluidic Device. Microsystems Nanoeng. 2020, 6 (1), 1–13. https://doi.org/10.1038/s41378-020-00220-3.

(47) Choi, C. H.; Lee, H.; Weitz, D. A. Rapid Patterning of PDMS Microfluidic Device Wettability Using Syringe-Vacuum-Induced Segmented Flow in Nonplanar Geometry. ACS Appl. Mater. Interfaces 2018, 10 (4), 3170–3174. https://doi.org/10.1021/acsami.7b17132.

(48) Trantidou, T.; Elani, Y.; Parsons, E.; Ces, O. Hydrophilic Surface Modification of PDMS for Droplet Microfluidics Using a Simple, Quick, and Robust Method via PVA Deposition. Microsystems Nanoeng. 2017, 3 (1), 16091. https://doi.org/10.1038/micronano.2016.91.

(49) Bauer, W. A. C.; Fischlechner, M.; Abell, C.; Huck, W. T. S. Hydrophilic PDMS Microchannels for High-Throughput Formation of Oil-in-Water Microdroplets and Water-in-Oil-in-Water Double Emulsions. Lab Chip 2010, 10 (14), 1814–1819. https://doi.org/10.1039/c004046k.

(50) Kim, J. Y.; Baek, J. Y.; Lee, K. A.; Lee, S. H. Automatic Aligning and Bonding System of PDMS Layer for the Fabrication of 3D Microfluidic Channels. Sensors Actuators, A Phys. 2005, 119 (2), 593–598. https://doi.org/10.1016/j.sna.2004.09.023.

(51) Deshpande, S.; Caspi, Y.; Meijering, A. E. C.; Dekker, C. Octanol-Assisted Liposome Assembly on Chip. Nat. Commun. 2016, 7 (1), 10447. https://doi.org/10.1038/ncomms10447.

(52) Kim, S. C.; Sukovich, D. J.; Abate, A. R. Patterning Microfluidic Device Wettability with Spatially-Controlled Plasma Oxidation. Lab Chip 2015, 15 (15), 3163–3169. https://doi.org/10.1039/C5LC00626K.

(53) Jahangiri, F.; Hakala, T.; Jokinen, V. Long-Term Hydrophilization of Polydimethylsiloxane (PDMS) for Capillary Filling Microfluidic Chips. Microfluid. Nanofluidics 2020, 24 (1). https://doi.org/10.1007/s10404-019-2302-2.

(54) Xia, Y.; Whitesides, G. M. SOFT LITHOGRAPHY. Annu. Rev. Mater. Sci 1998, 28, 153–184.

(55) Qin, D.; Xia, Y.; Whitesides, G. M. Soft Lithography for Micro- and Nanoscale Patterning. Nat. Protoc. 2010, 5 (3), 491–502. https://doi.org/10.1038/nprot.2009.234.

(56) Utada, A. S.; Chu, L.-Y.; Fernandez-Nieves, A.; Link, D. R.; Holtze, C.; Weitz, D. A. Dripping, Jetting, Drops, and Wetting: The Magic of Microfluidics. MRS Bull. 2007, 32 (09), 702–708. https://doi.org/10.1557/mrs2007.145.

(57) Kim, S.-H.; Kim, J. W.; Cho, J.-C.; Weitz, D. A. Double-Emulsion Drops with Ultra-Thin Shells for Capsule Templates. Lab Chip 2011, 11 (18), 3162–3166. https://doi.org/10.1039/C1LC20434C.

(58) Guo, J.; Hou, L.; Hou, J.; Yu, J.; Hu, Q. Generation of Ultra-Thin-Shell Microcapsules Using Osmolarity-Controlled Swelling Method. Micromachines 2020, 11 (4). https://doi.org/10.3390/MI11040444.

(59) Xu, S.; Nisisako, T. Polymer Capsules with Tunable Shell Thickness Synthesized via Janus-to-Core Shell Transition of Biphasic Droplets Produced in a Microfluidic Flow-Focusing Device. Sci. Rep. 2020, 10 (1), 4549. https://doi.org/10.1038/s41598-020-61641-8.

(60) Terekhov, S. S.; Smirnov, I. V; Stepanova, A. V; Bobik, T. V; Mokrushina, Y. A.; Ponomarenko, N. A.; Belogurov, A. A.; Rubtsova, M. P.; Kartseva, O. V; Gomzikova, M. O.; et al. Microfluidic Droplet Platform for Ultrahigh-Throughput Single-Cell Screening of Biodiversity. Proc. Natl. Acad. Sci. U. S. A. 2017, 114 (10), 2550–2555. https://doi.org/10.1073/pnas.1621226114.

(61) Vian, A.; Favrod, V.; Amstad, E. Reducing the Shell Thickness of Double Emulsions Using Microfluidics. Microfluid. Nanofluidics 2016, 20 (12), 1–9. https://doi.org/10.1007/s10404-016-1827-x.

(62) Vian, A.; Reuse, B.; Amstad, E. Scalable Production of Double Emulsion Drops with Thin Shells. Lab Chip 2018, 18 (13), 1936–1942. https://doi.org/10.1039/c8lc00282g.

(63) Kiss, M. M.; Ortoleva-Donnelly, L.; Beer, N. R.; Warner, J.; Bailey, C. G.; Colston, B. W.; Rothberg, J. M.; Link, D. R.; Leamon, J. H. High-Throughput Quantitative Polymerase Chain Reaction in Picoliter Droplets. Anal. Chem. 2008, 80 (23), 8975–8981. https://doi.org/10.1021/ac801276c.

(64) Fallah-Araghi, A.; Baret, J. C.; Ryckelynck, M.; Griffiths, A. D. A Completely in Vitro Ultrahigh-Throughput Droplet-Based Microfluidic Screening System for Protein Engineering and Directed Evolution. Lab Chip 2012, 12 (5), 882–891. https://doi.org/10.1039/c2lc21035e.

(65) Sun, C.; Liu, L.; Vasudevan, H. N.; Chang, K. C.; Abate, A. R. Accurate Bulk Quantitation of Droplet Digital Polymerase Chain Reaction. Anal. Chem. 2021, 93 (29), 9974–9979. https://doi.org/10.1021/acs.analchem.1c00877.

