## Supplemental Information for "Simplified, Shear Induced Generation of Double Emulsions for Robust Compartmentalization during Single Genome Analysis"

**Supplemental Figures:**

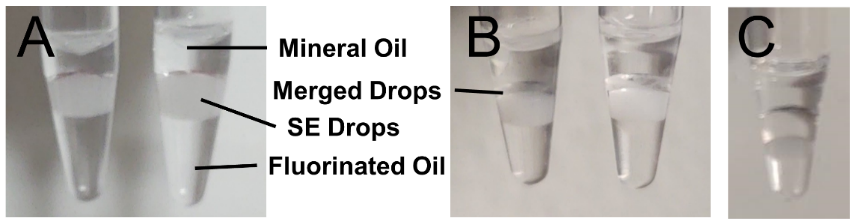

**Figure S1:** Surfactant lot and experimental conditions can result in variable merging in single emulsion drops during PCR. Replicate ddPCR experiments were performed using the same PCR reagents and surfactants. Tubes show minimal merging (A) and the formation of a merged layer after PCR (B). Using a different surfactant lot (C) significant drop merging was observed. SE drop merging is highly sensitive.

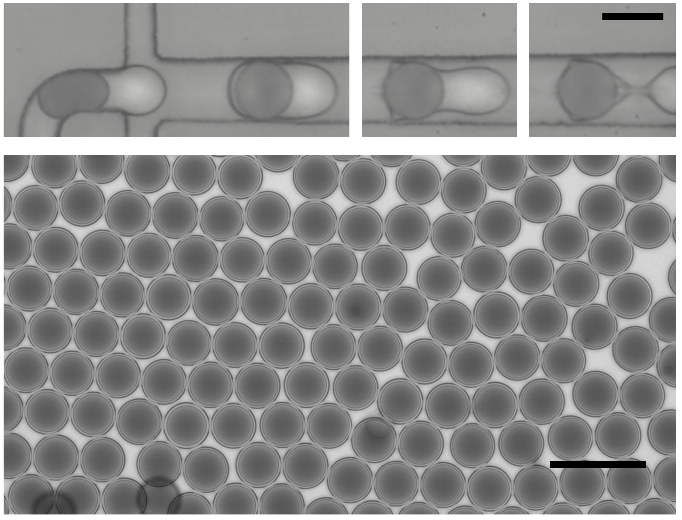

**Figure S2:** Figure S1. Monodisperse thin-shelled double emulsions formed by on-chip shearing. The scale bar is 100 μm.

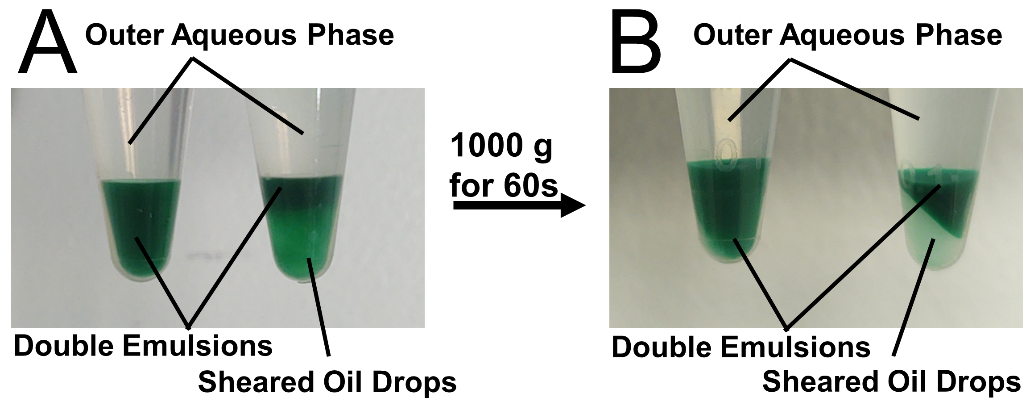

**Figure S3:** Figure S1. A) Left: Double emulsions formed without on-chip shearing. Right: Double emulsions formed with on-chip oil shearing (a mix of W/O/W DE and O/W SE drops). B) Mild centrifugation separates thin shell double emulsions from excess oil drops (Right). This centrifugation condition does not cause additional shearing of the oil shells (The left tube is unaffected). The outer aqueous phase remains clear indicating that the DE drops largely do not rupture.

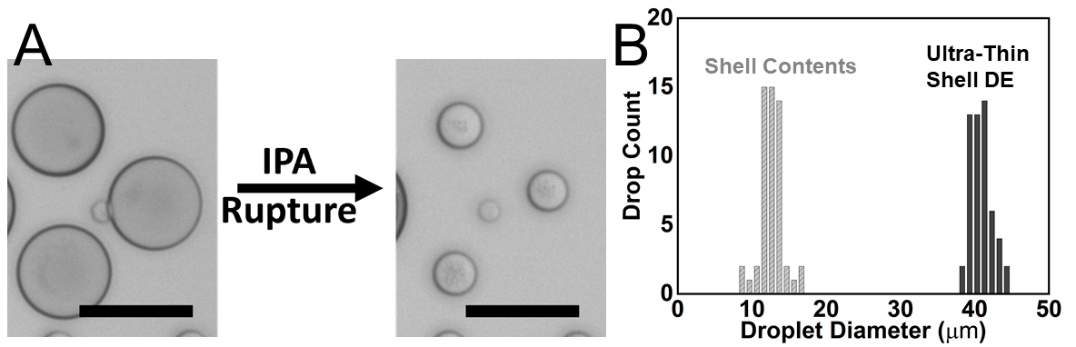

**Figure S4:** A) IPA induced DE rupture enables determination of shell thickness for very thing shell DE drops. The scale bar is 50 µm. B) Images are analyzed to obtain the total DE diameter before rupture and the oil drop diameter that results from the material contained in the shell.

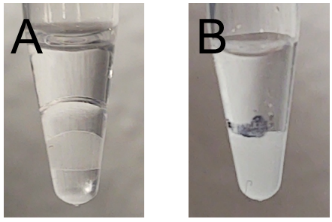

**Figure S5:** A) Large SE drops following PCR. A coalesced aqueous layer formed from merged drops can be seen below the mineral oil after thermal cycling. B) Large DE drops following PCR. The tube was marked to indicate the level of DE drops before thermal cycling. The majority of the DE survives PCR when the outer aqueous phase is osmotically matched to the aqueous core composition.

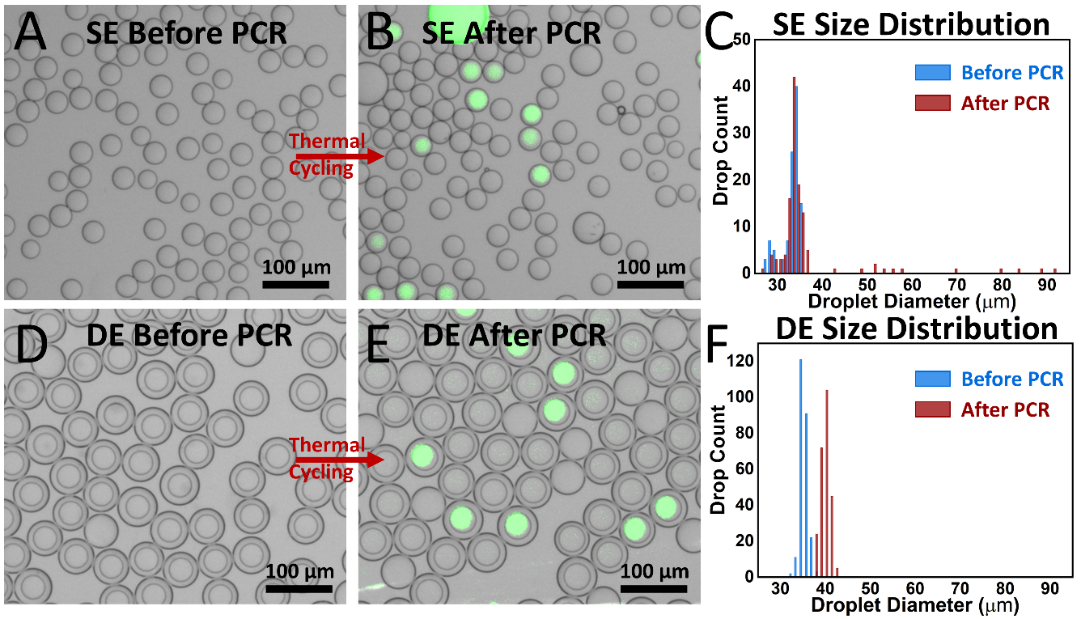

**Figure S6:** Comparison between ~40 µm single and double emulsion drops during ddPCR. A, D) Microscope images of drops after collection. B, E) Images of drops after 45 cycle PCR. SYBR green fluorescence indicates PCR amplification of SV-40. C, F) Histograms summarizing drop size distributions throughout the ddPCR workflow.

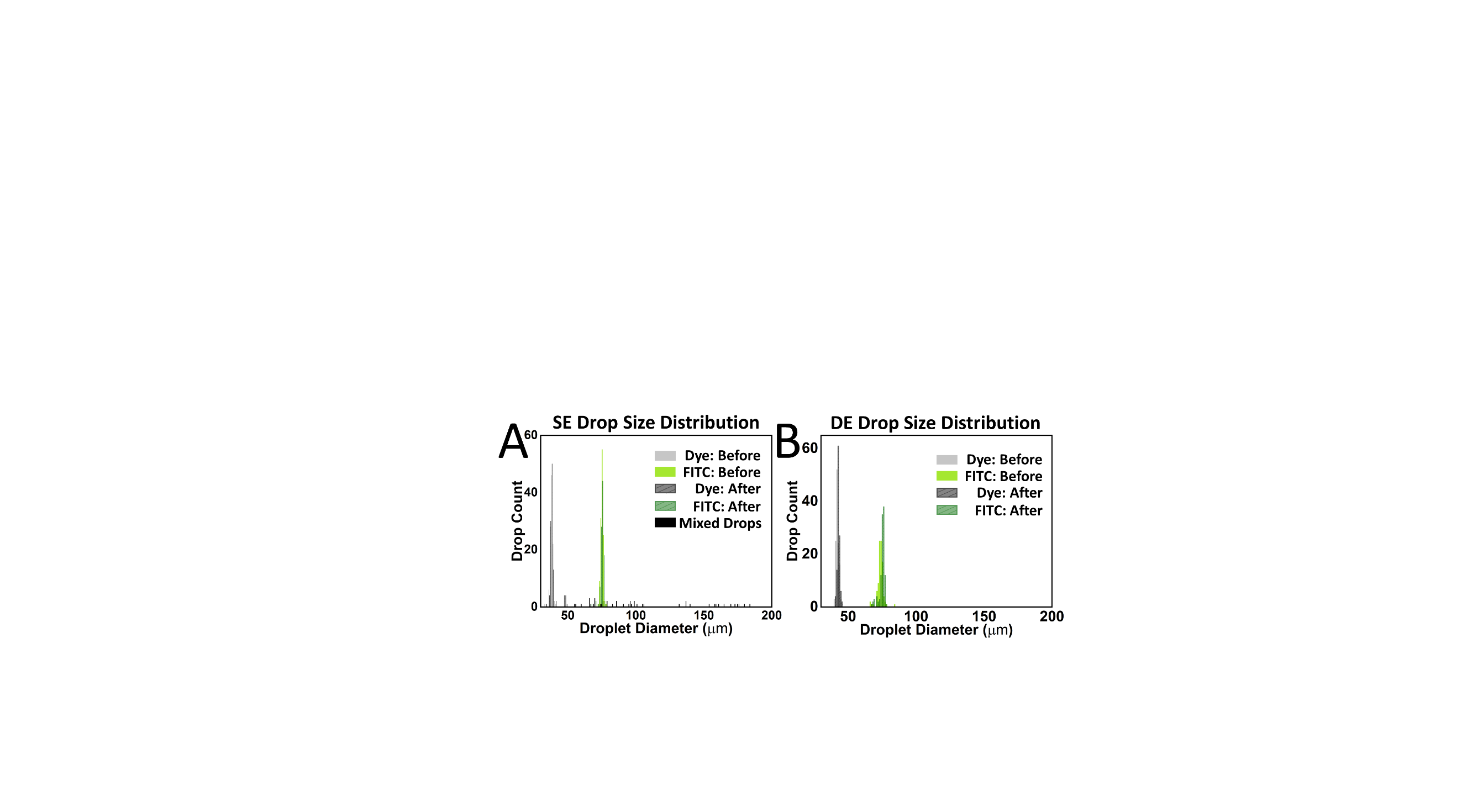

**Figure S7:** Histograms summarizing dop size distributions of heterogeneous mixed populations of dye and FITC drops. Single emulsion (A) and double emulsion (B) drops are analyzed before and after thermocycling. Heterogeneous double emulsion drops better maintain their size distribution as the SE shows larger merged drops following thermal cycling.

**Table S1.** Sequencing Primers

| Name | Sequence (5'-3') | Function |
| --- | --- | --- |
| BC | GCAGCTGGCGTAATAGCGAGTACAATCTGCTCTGATGCCGCATAGNNNNNNNNNNNNNNNGTTACGCGCCAGTAGAAACAAGG | Drop specific barcode with PCR amplification handles |
| R1-SV-F | ACACTCTTTCCCTACACGACGCTCTTCCGATCTCTCTCCAGACAAAGAACAACTGCC | SV-40 forward primer and Read 1 sequence |
| SV-R | CAGCGCTAGCTTCACCAACACCCTGCTCATCAAGAAG | SV-40 reverse primer and attachment handle to barcode |
| R1-λ-F | ACACTCTTTCCCTACACGACGCTCTTCCGATCTGGATAACACGCTCACCATGAAGC | λ forward primer and Read 1 sequence |
| λ-R | CAGCGCTAGCTTCAGTCACACTGTCAGGTGGCTC | λ reverse primer and attachment handle to barcode |
| BC-F | TGAAGCTAGCGCTGGCAGCTGGCGTAATAGCGAGT | Barcode forward primer and attachment handle to virus amplicons |
| R2-SE-BC-R | GTGACTGGAGTTCAGACGTGTGCTCTTCCGATCTACACCTTGTTTCTACTGGCGCGTAAC | Barcode reverse primer with single emulsion index and Read 2 sequence |
| R2-DE-BC-R | GTGACTGGAGTTCAGACGTGTGCTCTTCCGATCTGTTCCTTGTTTCTACTGGCGCGTAAC | Barcode reverse primer with double emulsion index and Read 2 sequence |
| P5-R1-F | AATGATACGGCGACCACCGAGATCTACACACACTCTTTCCCTACACGACGCTCTTCCGAT | P5 adapter attaches to Read 1 sequence |
| P7-R2-R | CAAGCAGAAGACGGCATACGAGATGTGACTGGAGTTCAGACGTGTGCTCTTCCGATCT | P7 adapter attaches to Read 2 sequence |

**Supplementary Video 1:** Oil shearing in a confined channel.

ASSOCIATED CONTENT

Supporting Information

The Supporting Information is available free of charge on the ACS Publications website.

Supporting Information (pdf)

Video of On-chip Shearing (mov)

AUTHOR INFORMATION

Corresponding Author

ORCID

Thomas W. Cowell: 0000-0002-7463-9339

Andrew A. Dobria:

Hee-Sun Han: 0000-0003-3616-291X

Author Contributions

T.W.C., and H.-S.H. have conceived of the idea and designed experiments. T.W.C, and A.D. designed and fabricated drop maker devices. T.W.C, and A.D. made drops, performed PCR, collected images, and analyzed data. T.W.C., and H.-S.H prepared the manuscript.
